# Assessment of different seedling production techniques of *Euterpe edulis*

**DOI:** 10.1101/2020.02.07.937755

**Authors:** Leonardo Lima Pereira Regnier, Maria Luiza Faria Salatino

## Abstract

*Euterpe edulis* is an endangered species with high importance ecologically and economically. Seedling production seems to be one of the most important alternatives to population recovery. Besides that, the knowledge of seedling production methods’ influence over germination is very restricted. Thus, this study aimed to evaluate the effects of parent populations, germination conditions, and the substrate to commercial seedling production of *E. edulis*. Nine thousand seven hundred and thirteen seeds were distributed between the heated water and control, greenhouse and open-field treatments. The parent population presented high differences between most of the germination indexes. Influencing the germination rate, mean germination time and germination speed, but not affecting synchrony and uncertainty indexes. Heated water treatment did not affect any of the studied indexes, presenting a close pattern of germination over time, indicating it is an appropriate method for seedling production. Greenhouse and open-field treatment presented variations at the same indexes affected in the parent population analysis. The most profitable method for *E. edulis* seed germination was the greenhouse production method, which provided the best indexes results.

## INTRODUCTION

*Euterpe edulis* Mart. also known as *Palmiteiro*, is one of the most representative and explored species from the Tropical Atlantic Rainforest [1]–[3], and is considered one of the most important [4]. Its slim, cylindrical, and straight stipe with a great green sheath is easily recognizable [1]. This species presents low and slow germination due to the mechanical structure of the seeds. These are recalcitrant; thus, they are not resistant to dry and/or temperature reduction. These features contribute to its long-term life cycle [5], [6]. Seedling growth seems to be limited by light exposure in natural conditions [7]. Adult individuals only reach sexual maturity between six and nine years old. These characteristics also hamper the natural regeneration of its populations.

This species also plays a relevant ecologic role as a food source for a great number of animals like birds, mammals, and insects [6], [8]. These interactions contribute to the reduced number of viable seeds that stay disposable to germinate and recover the population. An even lower number of germinated seeds reaches one-year-old [6].

The economic relevance of this species is its use as the heart of palm (the youngest part of the stipe, apical meristem, and young leaves) source. The heart of palm from this species presents high economic value and is an important product in the Brazilian and international markets [3], [4], [8], [9]. Besides, this species does not present tillers from the stipe basis [4]. Thus, harvesting the heart of palm from *E. edulis* implicates in the individual’s death. Furthermore, the illegal harvesting of this species heart of palm has been focusing on individuals at least 2 meters high. This criterion not only affects young individuals but also those in the reproductive phase (between six- and nine-year-old individuals), which is incompatible with this species slow life cycle, leading to a reduction in the natural populations and extinction [1], [4], [9].

*E. edulis* is also a key species in ecologic studies of the Atlantic rainforest [9]. Other authors have demonstrated that programs seeking *in situ* recovery must focus on seed banks’ maintenance and replanting techniques to effective forestry management [2], [5]. Thus, scientific research development in seedling production and ecophysiology is essential to guide recovery programs and/or commercial production [4], [5], [7], [10].

The demand for reproduction, quality, seed conservation, and seedling production technologies has been growing due to recovery programs and the recent inclusion of *E. edulis* species in landscaping projects [11]. Seedling production has also been pointed out as the main method for the management of natural populations of *E. edulis* [4], [12]. However, this knowledge is still very limited. Only recently, two basic aspects, how the removal of the mesocarp and light exposure influences over germination has been recognized [13]. The lack of technologies and knowledge in the exploration of native forestry resources [8] associated with natural habitat destruction are the main causes that still have been taken the native species to decline [8], [9].

In general, parent plants seem to influence the seedling quality and germination rate of its seeds [12], [14]. Most recently, to *E. edulis*, great diversity has been discovered and better explored [9], [15]–[17]. Besides that, the comprehension of how these differences between populations could affect germination parameters is not adequately known [9]. This aspect could provide important information in seedling production since the choice of a progenitor population used as seed sources could affect seedling production [12], [14].

The mechanical structure of this species seeds seems to hinder water penetration, generating great variations between seed germination rate [6]. One possible method to overcome this is by using hot water [18], [19], which also makes the removal of fruit pulp easier [5]. However, this method could reduce seed viability and reduce germination. Previous studies concerning the impact of hot water treatment on mineral bioaccessibility have already been conducted [20]. Nevertheless, possible germinations effects of this technique have not been extensively recognized in the literature.

In order to promote germination, in the commercial seedling production context, there are two main processes, open-field germination, where seeds are directly exposed to environmental conditions such as sunlight and temperature, or germination in a greenhouse, where seeds are exposed to greater temperature and moisture but less sunlight than the production area [21], [22]. These different techniques could also impact seed germination and seedling persistence [23], and aspects that have not yet been explored to *E. edulis*.

Therefore, due to the high ecological and economic importance associated with the great demand for reproductive techniques of this species. This study aimed to evaluate the effects of parent populations, heated water processing, and the open-field production approach to commercial seedling production of *E. edulis*.

## MATERIAL AND METHODS

This study was conducted at the Harry Blossfeld plant nursery of São Paulo, situated in Cotia (23°36’30.0”S 46°50’48.9”W). According to Köppen’s climate classification, the study region presents Cwa, the altitude tropical climate [24]. Featuring concentered rains during summer, dry winter, and the highest mean temperature above 22°C.

Plant material was collected in the western region of São Paulo in July and August 2018. Population I is 24 km distant from populations II and III, and these last two were 1 km away from each other. Fruits were gathered directly from the treetop and fallen fruits were also collected. All materials were kept in open plastic bags at room temperature for 2 days during the processing stage.

The study was divided into three steps. First, seeking to evaluate the parent population influence, approximately one thousand seeds from each of the three different populations were evenly distributed between two repetitions of the standard treatment [25]. This treatment consisted of exocarp and mesocarp removal with a sieve and running tap water. Seeds were planted in white trays containing vermiculite as a substrate and kept in a greenhouse with white plastic covering and a fogging watering system with periodic activation every 35 min.

The second part consisted of six thousand seeds, from population I, divided between two replicas of the 2 treatments. The control treatment used was the same as the standard treatment used in the first part of the study. While in the second treatment, all fruits were previously submerged in the heated water at 80 °C, seeking to facilitate the exocarp and mesocarp removal with a sieve and tap water. In sequence, seeds were planted in an open-field plant bed containing vermiculite as a substrate without any kind of covering and irrigation two times daily.

Seeking to evaluate seed development in open fields, 1 000 fruits from the same population, submitted to the standard treatment, and planted as described in the second part were sowed in plant beds containing vermiculite as substrate without any kind of covering and irrigation two times daily, 8 am and 4 pm.

Plant emergence was recorded 134 days after seeding, with measurements nearly every 7 days.

The evaluation consisted of the use of indexes as mean germination time, or mean length of incubation time [26], and the standard deviation was calculated as proposed by Haberlandt in 1875 [27]. All other indexes were calculated as presented by Lozano-Isla *et al.* [28].

All data were compiled into the Excel component of the Microsoft Corporation Office pack and represented through graphics and tables. The most common germination indices were obtained using GerminaQuant software [28]. All statistical tests were executed with individual data of the replicas and performed using GerminaQuant, based on R software [29]. The results were tested by ANOVA and in sequence submitted to the Tukey test, adopting a critical p-value of 1% (α < 0.01).

## RESULTS AND DISCUSSION

### Parent population

The parent population seems to influence the Germination proportion (GRP), Mean germination rate (MGR), and Germination speed (GSP) (Fig. 1A, 1B, 1D). While Uncertainty (UNC) and Synchrony (SYN) indexes were not different (Fig. 1E, 1F). Populations I and II presented the highest germination rates (91.3% and 78.57%) and were consistently different from population III (22.9%). The mean germination rate index (MGR) indicates, in this study, the frequency of germinations, as the mean germinations/day. Thus, it is the reciprocal information of mean germination time (MGT) [26].

**Figure 1.**
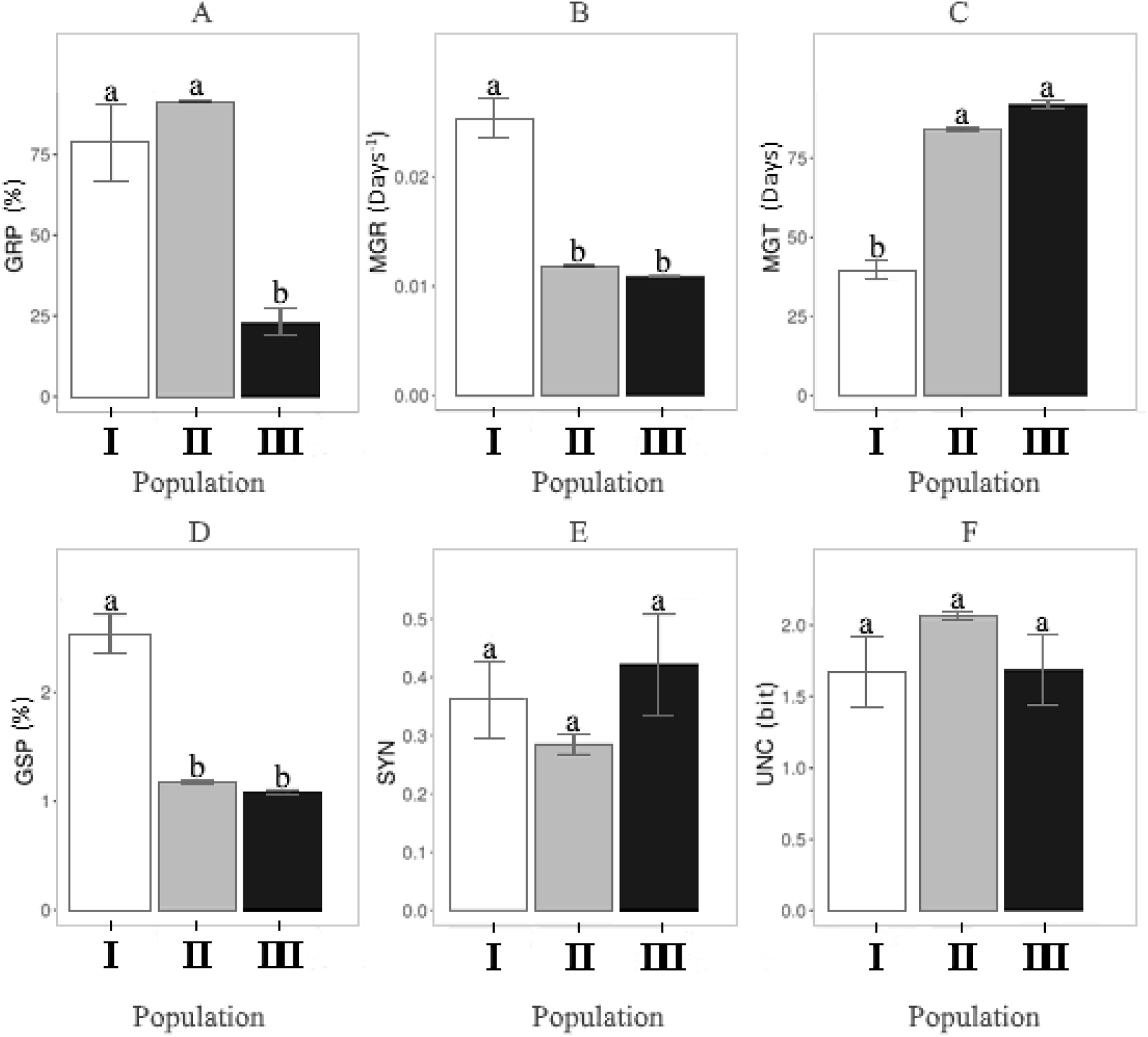
*E. edulis* germination indexes according to seed population origin **(I, II, III). A** - Germination Proportion (GRP), **B** - Mean germination rate (MGR), **C** - Mean germination time (MGT), **D** - Germination speed (GSP), **E** - Synchronization index (SYN), and **F** - Uncertainty index (UNC). Lowercase letters present statistical differences using p<0.01. Error bars represent the standard error mean.

The mean germination time (MGT) or mean length of germination time demonstrates the required time for one seed of some species to germinate [26], [30]. It is an important index insofar as it demonstrates the mean expected time for most seeds of a seed lot to germinate. Therefore, it is possible to estimate the necessary time for some specific culture to grow properly. The data between the populations were significantly different (Fig. 1). Only populations II and III presented similar mean germination rates (84.36 and 91.9, respectively). Population I reached stabilization about half the time obtained by the other populations, and also greater germination speed in this study (Fig. 1D). These are a valuable aspect of the seedling production context. Thus, population I has the most profitable offspring, seeking better production.

Mean germination time is associated with the general characteristics of the seed species, seed lot quality, and environmental conditions [5], [31]. The unbridled harvesting of adult individuals that led *E. edulis* natural population decline, related to the reproduction between related individuals, could have led some populations to less vigorous offspring, a process called endogamy depression [12], which directly affects seed quality [2], [14]. This species presents an aggregated spatial distribution of seedlings, favoring intraspecific competition and greater vulnerability to diseases and plagues [32]. These features, associated with short distance gene flow [14], could result in high endogamy, reducing seed diversity and quality. This process directly impacts the seed harvesting approach seeking conservation projects [14], since the great variation between seed lots hampers commercial seedling production [12]. The studied parental populations had considerable variations in germination rate and also germination speed indices (Fig. 1). Indicating that choosing some population offspring not only influences the exploitation of a seed lot, based on the germination proportion, but also on the time required for seedling emergence. Thus, the process of choosing a parental population for seedling production seems to be a very important aspect to be considered.

The germination rates of the studied parent populations provided significant differences between population III and the others (Fig. 1A). The cumulative germination course also reveals the discrepant performance of this population compared to the others (Fig. 2). Germination of *E. edulis* seems to differ greatly due to genetic variations between fruits, even when they are harvested at the same time and present the same developmental stages [12]. An important aspect of genetic variability in the reproductive system [14]. It is estimated that the gene flow between individuals of *E. edulis* is 56 meters using DNA markers [14], which favors genetic isolation between populations and high variations between them [9]. However, natural population decline seems to have led to endogamy depression. Parent plant age also seems to influence the germination indices [3], [6], and the reproductive season [2]. Germination presented in other studies comprehends the range of 16% to 52% at 12 days after sowing [6] and from 44% to 73% between 100 and 150 days after seeding [12], [33]. The present study data indicate a higher germination rate in populations I and II than those previously mentioned values found by those authors (Fig. 2). Besides that, none of the studied populations reached at least 16% of germination during the first 12 days after sowing, as mentioned by Bovi [6]. However, these discrepancies are common between non-domesticated species [14] due to genetic variations, and also the environmental conditions related to the locations were the studies were performed.

**Figure 2.**
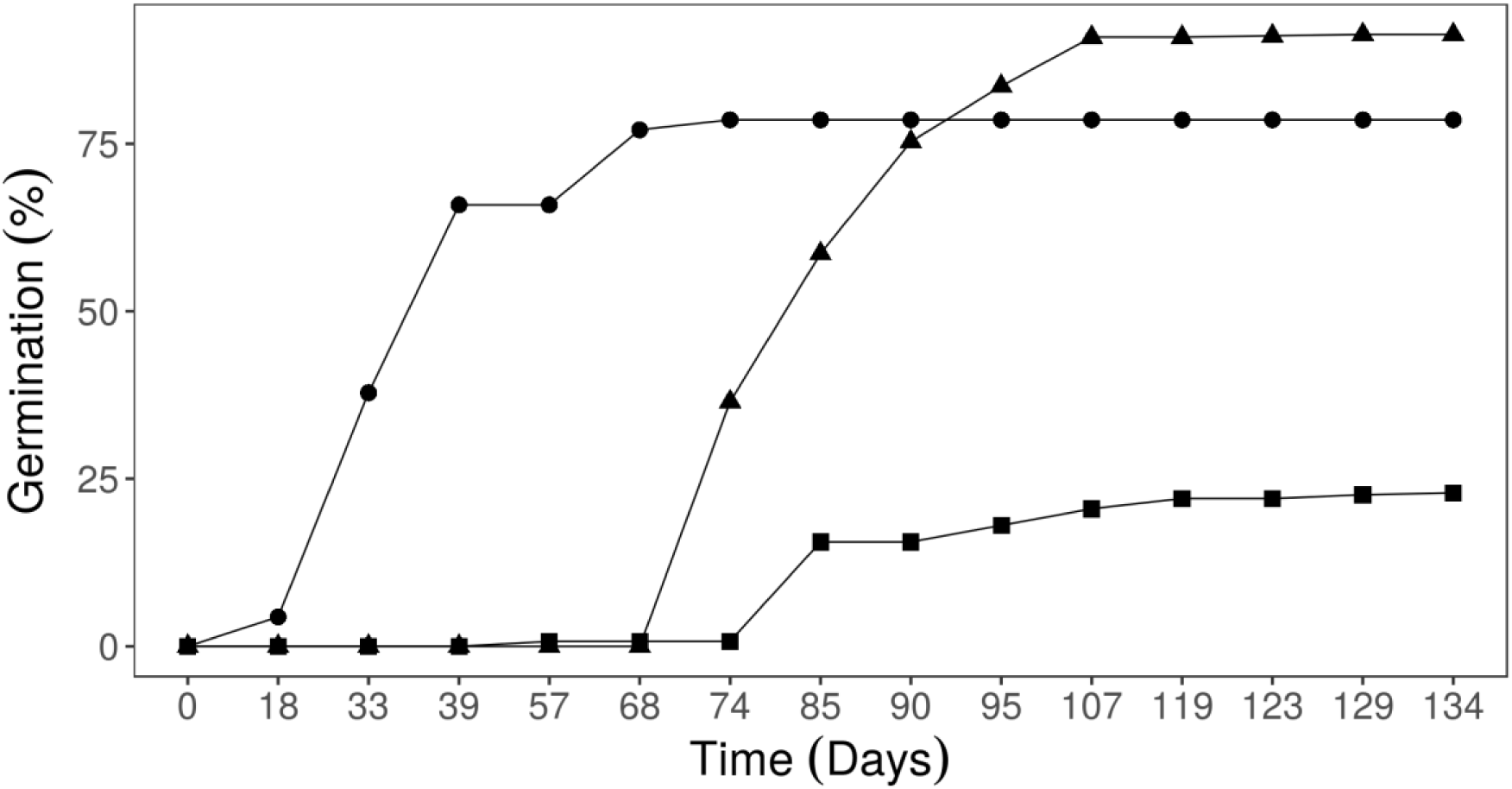
Cumulative germination of *E. edulis* according to seed origin: Population I (●), Population II (▲), and Population III (■).

It is important to note that the germination pattern was very different between the populations (Fig. 2). Although populations I and II reached relatively close germination results at the end of the study period, their trajectory until reach those results were not equal. Most recent studies have shown that, in general, diversity between this species’ populations remains high considering fragmentation, habitat destruction, and selective exploration [9]. Our results also presented a great diversity of germination responses between the populations. Corroborating this information, conspicuous variations between germination indexes were observed when comparing populations of *E. edulis*.

In commercial seedling production, understanding the factors that could impact seed lot quality is very important, especially for understanding suboptimal results [34]. These study data indicate that choosing the parental population as a seed lot provider could drastically modify seed lot quality, being a crucial aspect in the seedling production context.

Non-cumulative germination pattern indicates that, in general, germinations are concentrated at some specific period, when most of the seeds geminate synchronically (Fig. 3). Besides that, the values of the germination rate reached by each population varies. The population III pattern is similar to other populations but delayed and with lower values (Fig. 3). Non-cumulative germination analysis could provide some important information. In this study, seeds from different populations seem to exhibit a similar pattern, with relatively concentrated germinations at some specific time, diverging only by when this higher emergence period happens. This fact explains why the uncertainty and synchrony indexes were similar between the studied populations (Fig. 1). Insofar as they presented a very similar distribution pattern but with contrasts about the time required to reach great germination occurrence. The different parent populations seem to provide variations on when germination occurs and the relative distribution of germination all over time.

**Figure 3.**
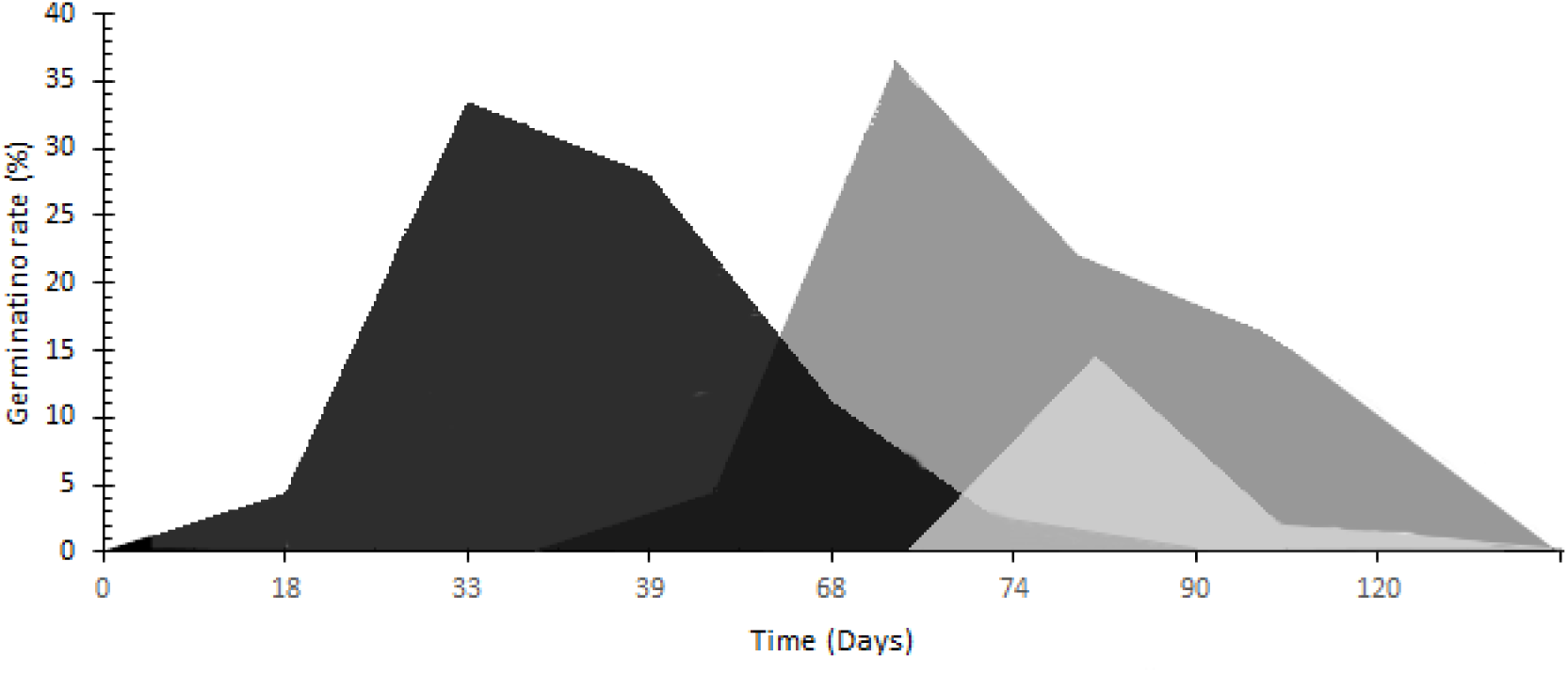
Non-cumulative germination of *E. edulis* according to seed origin: Population I (■), Population II (■), and Population III (■).

Although, it is important to note the long-term germinations presented by this species. Previously, other authors have already mentioned that the greater variations in the required time to seedling emergence is an adaptive strategy required to maintain a proper seed bank [12]. An essential implication of ecologic dynamics because if all the seeds from a single cohort present unsynchronized and long-term emergence, this single reproduction event could provide recruits to the population during very long periods, even with dramatic reductions in the parental population. However, these features hamper seedling production [35] because this context only favors faster and synchronized germinations.

### Heated water treatment

There was no difference between the control and heated water treatments. All the analyzed indexes did not present significant variations. It seems that using heated water only provides a slight reduction in germination proportion and mean germination time (Fig 4). In general, heated water promotes mechanical ruptures that can easily lead water to permeate [36]. However, high temperatures could also kill the embryos, reducing seed viability [5]. Our results show that any of the studied indexes related to the time required for germination, speed, or germination rate were substantially affected by this method at this study parameters. Indicating that there is no significant reduction in seed viability when exposed to this temperature. However, if *E. edulis* presents a mechanical structure that hinders water penetration, as mentioned by other authors [6], heated water at 80°C does not provide its rupture. Better anatomic and germination studies are required to clarify the seed structure and provide methods to improve the germination rates of *E. edulis*. Cursi & Cicero [5] found that *E. edulis* fruits, when immersed during 20 minutes on water at 40°C, presented higher germination. While exposure to 55°C water before processing seems to be harmful to embryos. Our results conflict with this information. However, the consistent variations observed in germination indexes provided by seeds’ origin, and environmental conditions adopted during the experiment could also be responsible for this kind of discrepancies between studies.

**Figure 4.**
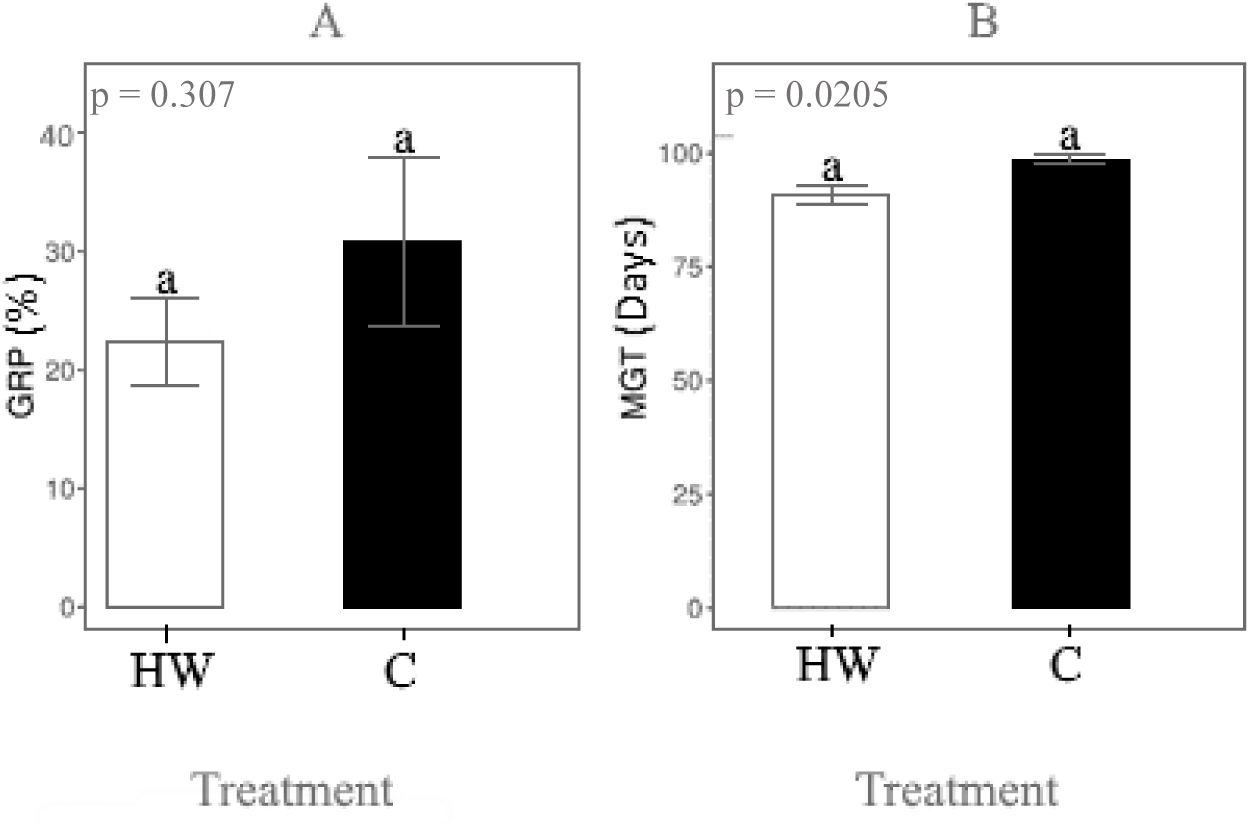
*E. edulis* germination indexes according to seed processing treatments: Heated water (HW) and control (C). **A** - Germination Proportion (GRP), **B** - Mean germination time (MGT). Lowercase letters represent statistical differences adopting p<0.01. Error bars present the standard error mean.

The cumulative germination pattern was also very similar between the treatments (Figure 5), emphasizing that this procedure does not substantially modify seed germination. Besides that, adopting a heated water procedure makes the removal of fruit external parts easier, and it seems to not substantially affect the seedling production seeking commercial proposes, which was also mentioned in previous studies [5]. Therefore, this kind of procedure could be used in the seedling production context, without major harmful effects.

**Figure 5.**
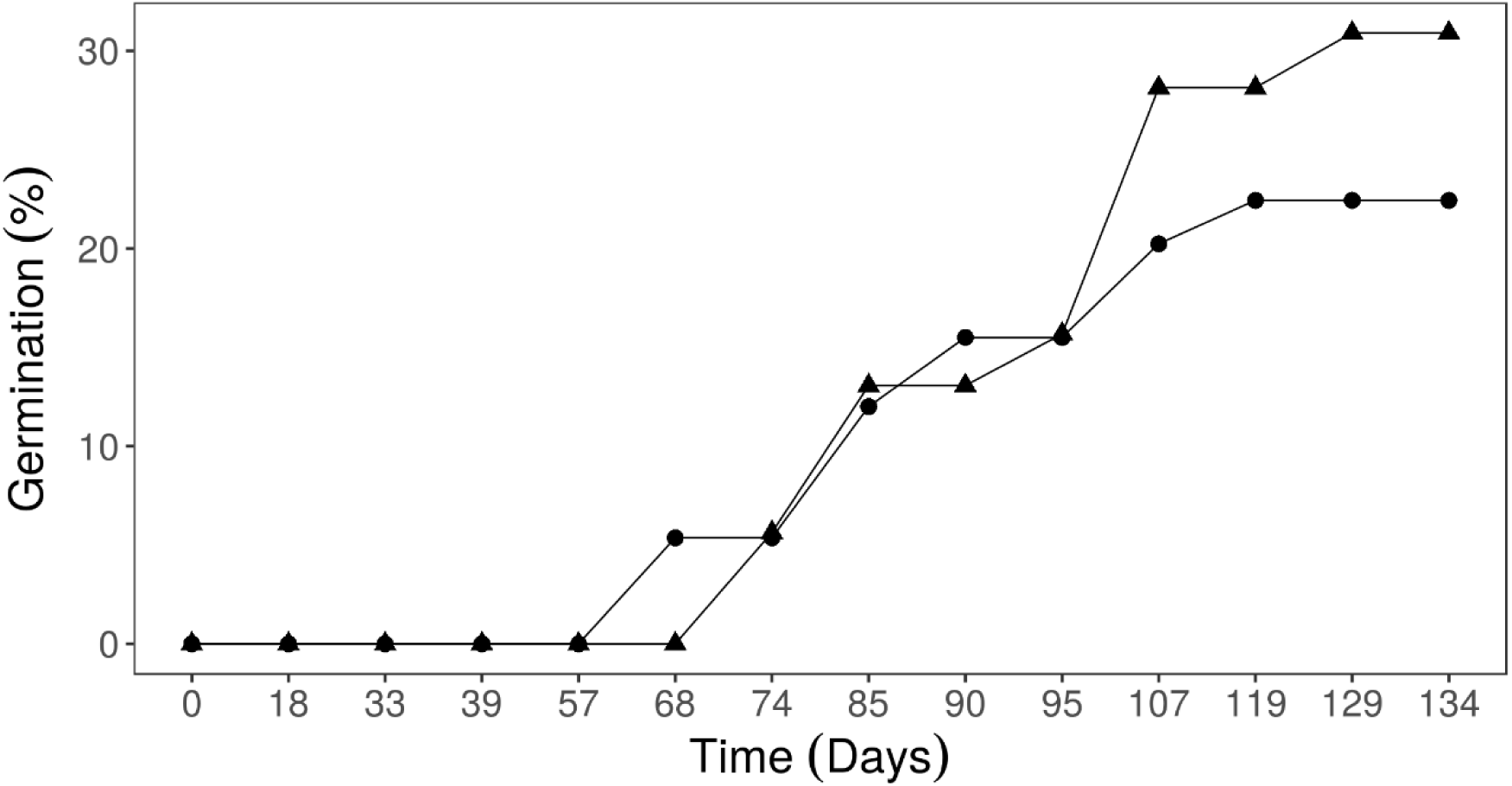
Cumulative germination of *E. edulis* according to seed processing. Heated water (●) and Control group (▲).

### Open-field and greenhouse production process

Analyzing the production process of open field and greenhouse, some variations were significant to all the analyzed parameters, except for the uncertainty and synchronization index (Fig. 6). The greenhouse process provided better results at all indexes, the required time to germination and stabilization were faster, and the germination rate was significantly greater, providing better exploitation of a seed lot. The greenhouse production process has been recognized to notably optimize plant production and reduce the limits of growth [37]. The results also emphasize that the greenhouse promotes faster and better germination of *E. edulis* seeds. Greenhouses usually promote a stable temperature, controlled moisture, and less susceptibility to environmental conditions. This group of features favors seedling emergence. Contrasting to the open field, the greenhouse promoted both better germination rate and reduced time required for germination, favorable aspects seeking commercial seedling production.

**Figure 6.**
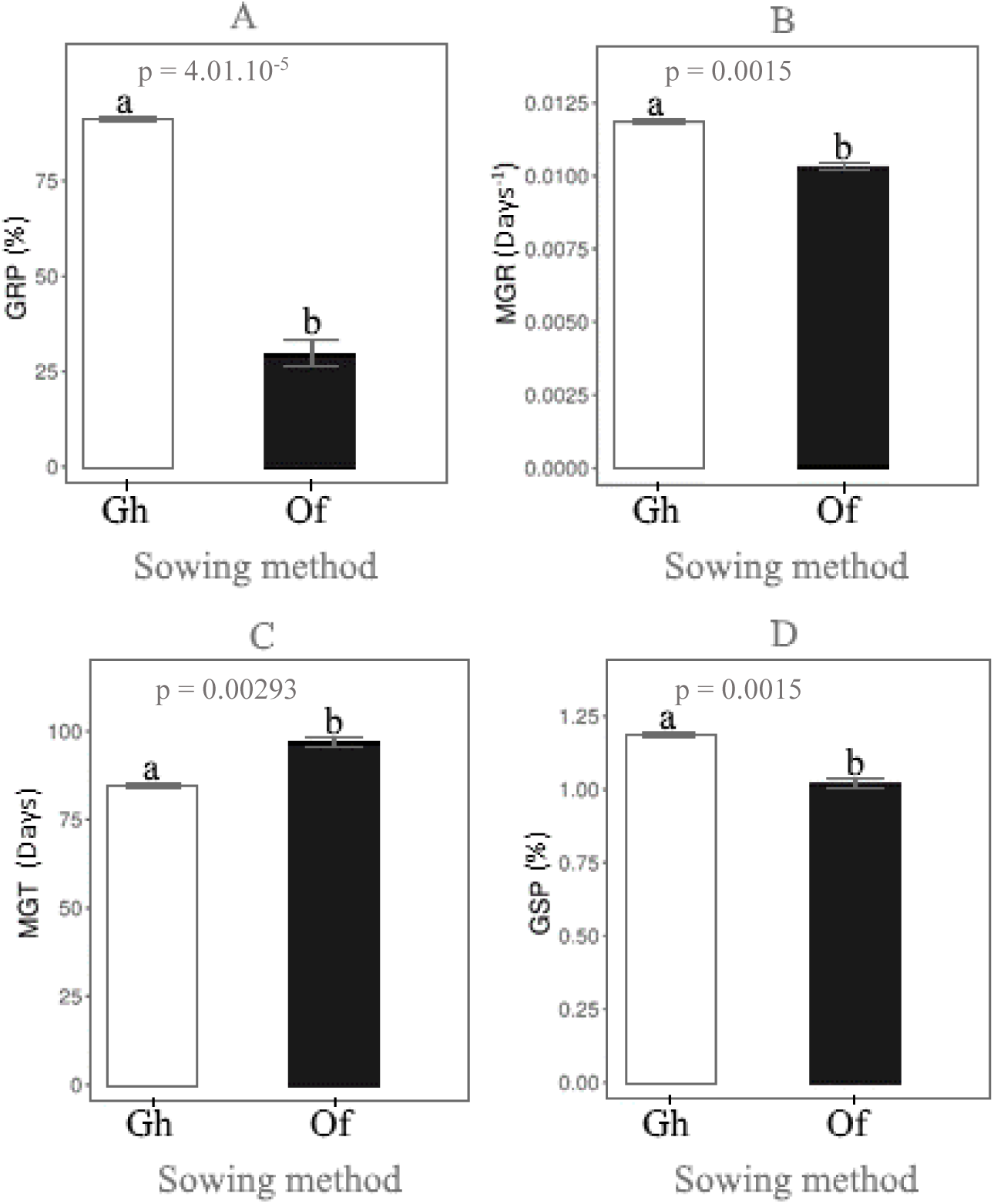
*E. edulis* germination indexes according to different sowing methods, greenhouse (Gh), and Open-field (Of). **A** - Germination Proportion (GRP), **B** - Mean germination rate (MGR), **C** - Mean germination time (MGT), and **D** - Germination speed (GSP). Lowercase letters represent statistical differences adopting p<0.01. Error bars present the standard error mean.

The cumulative seed germination pattern was substantially distinct (Fig. 7). It is already recognized that controlled conditions favor better expression of the germinative capacity of seed lots, while in the open field, the attack of animals and also microorganisms, reduces the viability of seeds affecting germination and seedling development [2], [38]. Open-field also propitiates higher sunlight exposure and less water provision. Water availability seems to be a crucial aspect of *E. edulis* seed development. Earlier life stages are the least tolerant to water deficit, and the resistance seems to gradually increase with age [10]. Thus, the lesser water provision in open-field treatment could be the main factor to reduce germination performance. In an ecological approach, open-field treatment also provides environmental conditions relatively closer to natural or anthropic disturbed areas [23]. Providing a set of environmental conditions that do not favor seedling establishment. Reduced performance of germination the open-field technique has also been reported in other species from Fabaceae [23]. Besides that, some authors affirm that even when this species is more exposed to environmental conditions, *E. edulis* plays the role of pioneer species, being the first mesophytic species to conquer disturbed environments [8].

**Figure 7.**
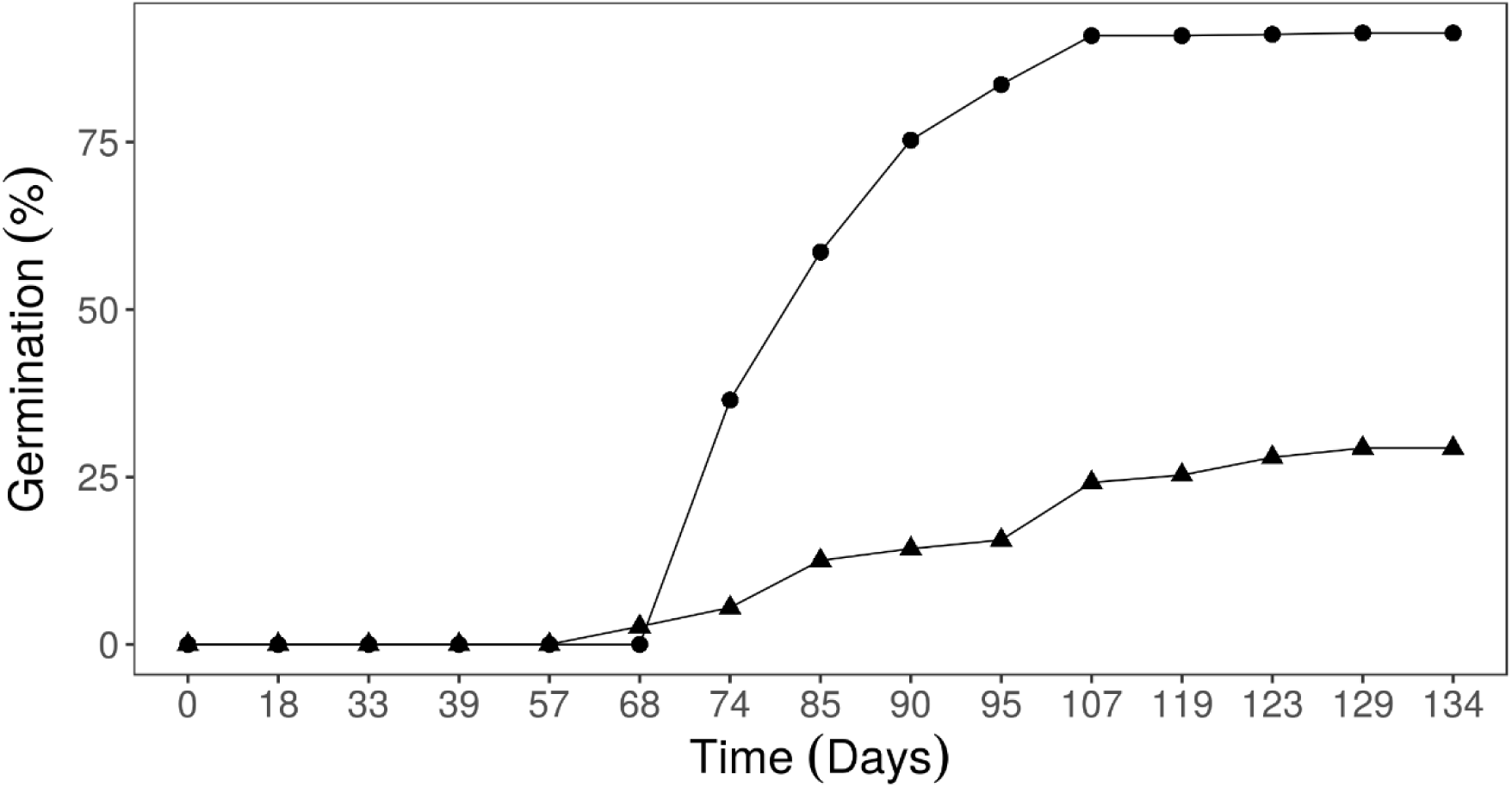
Cumulative germination of *E. edulis* according to the seed production process: Greenhouse (●) and Open-field (▲) production methods.

Even though the open-field production method does not provide the best environment for germination, this method is adopted for some crops because it requires less electricity and labor when compared to the greenhouse method [39], and facilities maintenance is cheaper than greenhouses, reducing the production cost [23]. Sometimes this practice is also used to recover degraded areas [38]. However, the cost-benefit ratio needs proper evaluations for further studies.

## CONCLUSION

The different parent populations seem to influence the germination rate, mean time to germination, and germination speed, but did not affect the uncertainty and synchrony indexes.

Water heated at 80°C did not affect germination indices, indicating that it is a feasible method for seedling production. The most profitable method for *E. edulis* seed germination is the greenhouse production method since it provides the best results, associated to germination, speed, and time required for germination.

## ACKNOWLEDGMENTS

I would like to appreciate Secretaria do Verde e Meio Ambiente of São Paulo due to its trainee programs. In addition, the workgroup of Harry Blossfeld. Also Rafaela C. Perez, Juliana de Lemos, and Caio G. Tavares Rosa, due to their consistent scientific support and encouragement.

